# A continental scale analysis reveals widespread root bimodality

**DOI:** 10.1101/2022.09.14.507823

**Authors:** Mingzhen Lu, Sili Wang, Avni Malhotra, Shersingh Joseph Tumber-Dávila, Samantha Weintraub-Leff, Luke McCormack, Xingchen Tony Wang, Robert B. Jackson

## Abstract

Recent studies of plant fine roots have greatly advanced our understanding of their geometric properties and symbiotic relationships, but knowledge of how these roots are spatially distributed across the soil matrix lags far behind. An improved understanding of broad-scale variability in root vertical distribution is critical for understanding plant-soil-atmosphere interactions and their influence on the land carbon sink. Here we analyze a continental-scale dataset of plant roots reaching 2-meters depth, spanning 19 ecoclimatic domains ranging from Alaskan tundra to Puerto Rican neotropical forest. Contrary to the common expectation that fine root abundance decays exponentially with increasing soil depth, we found surprising root bimodality at ~20% of 44 field sites —a secondary peak of fine root biomass far beneath the soil surface. All of the secondary root peaks were observed deeper than 60cm (with 33% below 1m), far deeper than the sampling depth commonly used in ecosystem studies and forestry surveys. We demonstrate that root bimodality is more likely in places with relatively low total fine root biomass, and is more frequently associated with shrubland vegetation but less with grassland. Further statistical analyses revealed that the secondary peak of root biomass coincided with unexpected high soil nitrogen contents at depth. By linking roots and nutrient distributions, we further demonstrate that deep soil nutrients tend to be underexploited by plant rooting systems, yet root bimodality offers a unique mechanism by which fine roots can tap into soil resources in the deep. Our findings suggest that empirical practices have often systematically overlooked root dynamics in deep soils, and as a result the current-generation global climate and vegetation models have relied on overly simplistic assumptions for plant rooting distribution.

## Main text

Terrestrial ecosystems face a pressing challenge due to increasing atmospheric CO_2_ concentration: how can land plants satisfy their increasing nutrient demand^1–3^ and consequently sustain the terrestrial carbon sink^4–7^? A central piece to this puzzle is the ecology of plant roots. Plant roots not only are key to the uptake of growth-limiting soil resources^8^, but are also central to soil carbon input through root turnover^9–12^ and carbon loss through priming^13^.

Recent studies have greatly improved our understanding of plant roots through the study of their “traits”, ranging from morphological traits such as root diameter^14–17^ to root-symbiont relationships such as mycorrhizal colonization rate^18,19^, in part thanks to the rise of publicly available harmonized root trait data^20–23^. However, these trait data mostly reflect micro-scale properties of individual root segments (*e.g.*, root diameter, root nitrogen concentration). A rooting-system level understanding of how these roots are distributed throughout the soil matrix still lags far behind (but see ref^24^), limiting our ability to scale up local, trait-based measurements to ecosystem-scale properties.

In particular, the ecology of deep fine roots (> 1m depth) is perhaps the least understood component of root ecology due to the difficulty in sampling them^25–29^. Previous studies have reported the importance of deep fine roots in aiding plants to take up deep water^30^, tap into deep soil nutrient reservoirs^31^ and convert photosynthetic carbon into deep soil organic matter with high efficiency^32^. Yet, despite the importance of deep fine roots, our decades of understanding of plant rooting systems has been mostly confined to the top 5-100 cm^33^ due to the paucity of deeper fine root data^22,34^.

The recently established National Ecological Observatory Network (NEON hereafter) uniquely addresses this data gap^35^. First and most importantly, the NEON soil megapit data (Methods) reaches an impressive depth of 2 meters below the surface, allowing us to examine fine root dynamics to an unprecedented depth (Fig.S1,Table.S1). Second, the NEON megapit root data are coupled with data on soil nutrients, soil physical properties, and soil moisture, making it possible to link root distributions to various abiotic factors. Third, the same standardized sampling approach is repeated across 44 sites ranging from Alaskan tundra to Puerto Rican neotropical forest^36^, enabling us to perform continental scale analysis across a range of ecosystem types and climate zones.

Leveraging these unique strengths of the NEON dataset, we evaluated three questions regarding the ecology of deep roots (1) How does the abundance of fine roots change with depth? (2) What are the biotic and abiotic factors that impact the distribution of fine roots with depth? (3) Are nutrients in deeper soils equally, under-, or over-exploited by fine roots compared with surface soil?

## Results and discussion

### Deep sampling reveals unexpectedly widespread root bimodality

We first examined the distribution of fine-root biomass along soil profiles up to 2m depth. Unexpectedly, we frequently observed rather unusual bimodal distributions of fine-root abundance (*i.e.,* dry root biomass per volume), which differs from the default expectation of exponential-decay distribution^37,38^. In Fig.1, we show the fine-root distribution at site SCBI (Smith Conservation Biology Institute NEON) as an example of root bimodality, in contrast to the typical unimodal exponential-decay at site YELL (Yellowstone National Park NEON). The bimodal SCBI site’s secondary peak of root biomass is found well beneath the soil surface (~1.3m, Table.S1), and well beyond where most sampling efforts stop^22,34^.

**Figure 1.**
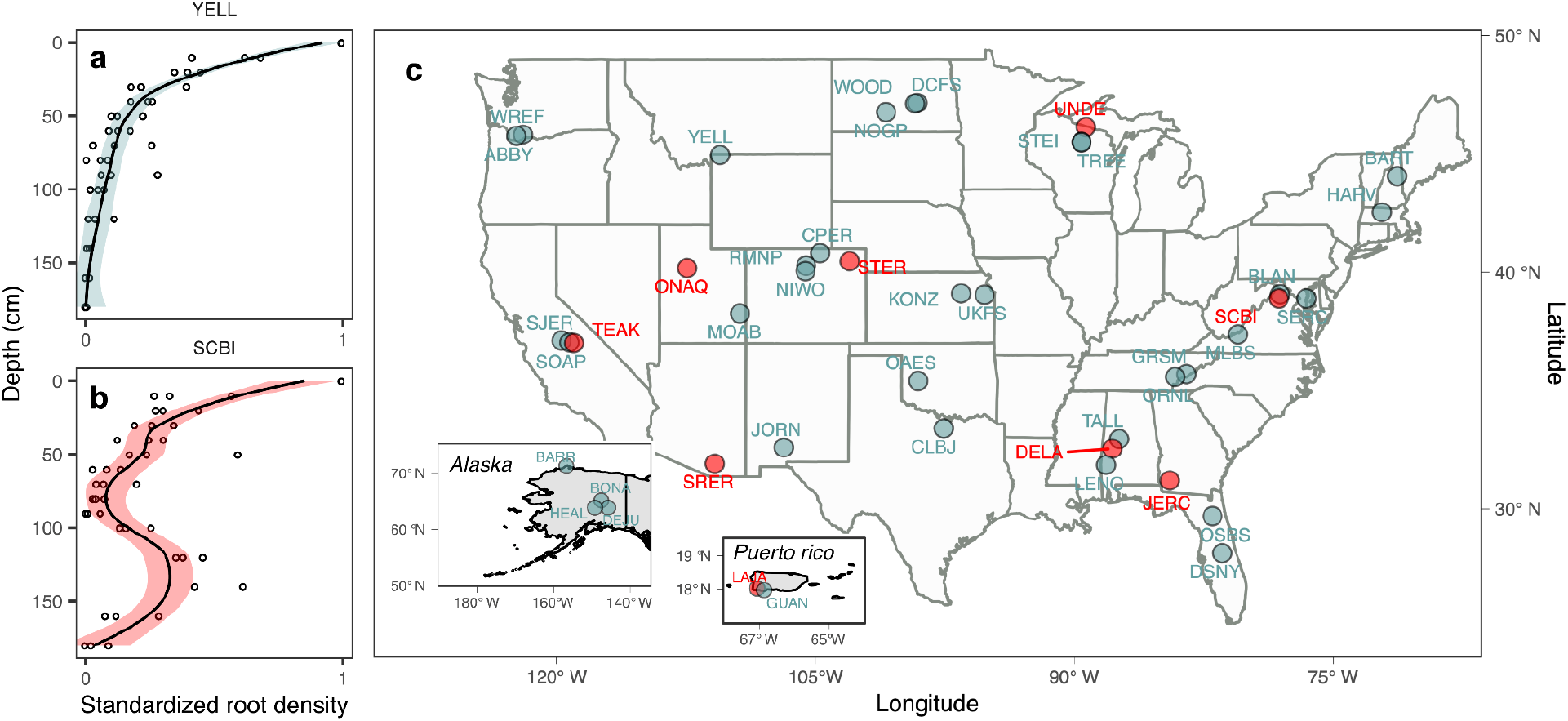
Bimodal rooting distribution is widespread. **(a)** An example depth profile (YELL) of the classical exponential decay of root abundance with increasing depth. **(b)** An example depth profile (SCBI) of a bimodal rooting distribution shows a secondary peak of root biomass far beneath the soil surface. Note that the root abundance is standardized based on the maximum abundance to facilitate visualization and cross-site comparison. Ribbons represent confidence intervals around LOESS fits (solid black), with red fill denoting bimodal distribution and blue denoting the unimodal exponential decay. **(c)** The spatial distribution of rooting profiles across 44 NEON sites. Detailed site descriptions can be found in Supplementary Information Table.S1 along with all 44 rooting profiles (Fig. S2).

What’s more, this surprising phenomena of bimodal rooting distribution is widespread across the continental US, displaying no obvious patterning (Fig.1c; bimodal sites in red, unimodal in blue). We estimate that nine out of 44 (~20%) sites in the NEON dataset have bimodal root distributions (see Methods). It is especially noteworthy that all 9 of these bimodal sites feature a secondary root peak deeper than 60 cm (with 33% below 1m; Table. S1), meaning these sites would NOT have been classified as bimodal using more common, shallower sampling depths.

As a natural phenomena, bimodality has been widely spotted across multiple science disciplines: from the bimodal body mass of African weaver ants^39^ to the bimodal mycorrhizal association in tropical rainforest^18^, from the distribution of water vapor in Earth’s atmospheric system^40^, all the way to the color and shape distribution of galaxies in our cosmos^41^. The observation of bimodality often indicates the coexistence of two contrasting regimes/processes, of which the underlying mechanism is unaccounted for, raising the question of what process can give rise to the observed root bimodality.

Intuitively, if there exists a singular mechanism underlying all these bimodal root distributions (9 out of 44 NEON sites), there’s a good chance we can detect the mechanism using statistical tools. In reality however, many processes can potentially cause bimodality of fine roots. For instance, the presence of a buried soil horizon rich in nutrients, surface drought coupled with presence of water supply in deeper soil, the presence of certain species that specialize in foraging for deep soil resources, to name a few. The potential multi-cause of bimodality might limit our ability to derive an universal explaining mechanism. But failure to pin down a singular cause would then suggest that different NEON sites might operate under very different rules and bimodality might be due to distinct site-specific mechanisms. In the next two sections, we evaluate how root bimodality is linked to a range of factors, across and within the NEON sites.

### Bimodality linked to site-level factors

We first analyzed whether the presence of root bimodality is linked to site-level features across all NEON sites. We built a classification model that takes into account 12 site-level features (Methods, Fig.2a) to examine the predictive power of each feature for bimodality. We chose a random forest algorithm for our classification model because (1) it is robust to the presence of outliers and nonlinear relationships and (2) it has a built-in validation mechanism that is particularly valuable for our small sample size (44 NEON sites). The resulting best performing classification model achieved an overall accuracy of 64% (correctly predicting the modality of 28 out of 44 sites), with a 66.7% accuracy for predicting bimodal and 63% accuracy for predicting unimodal sites. The importance ranking of features (Fig.2**a**) from this classification model identifies total root biomass (g/m2; per square meter surface to maximal pit depth) as the most important predictor of bimodality, followed by gross primary production (GPP), and a range of other features.

**Figure 2.**
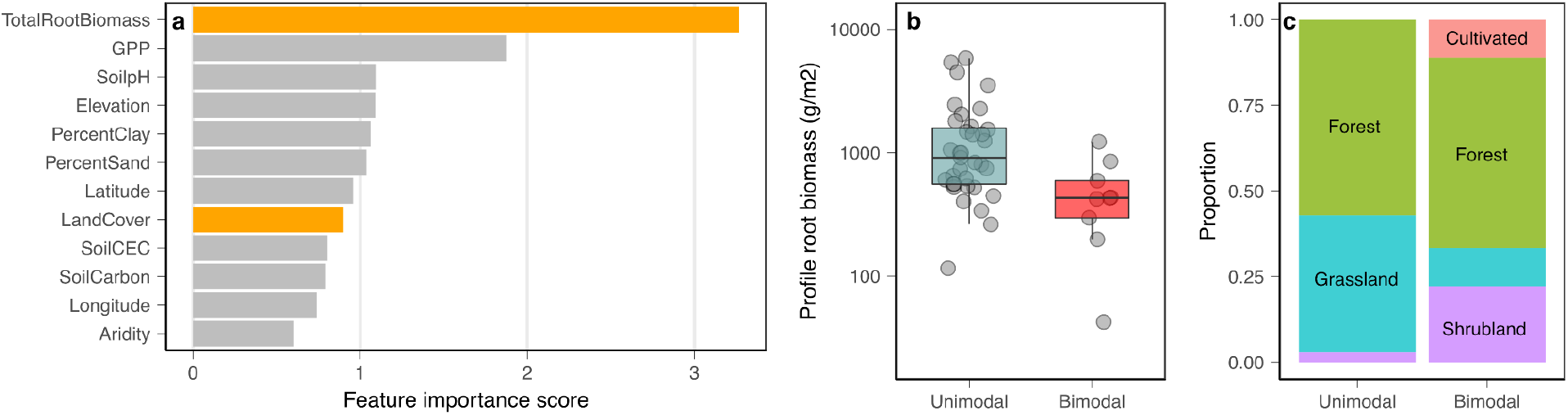
Root bimodality predicted by site-level factors. **(a)** Feature importance ranking from a random forest classification model (Methods) identify total root biomass (g/m2) as the most important factor in predicting the modality of a site. Of these 12 factors, land cover is the only categorical variable, the rest are continuous. **(b)** Unimodal sites have nearly 3 times more root biomass than that of bimodal sites (990 *vs.* 368 g/m2; *p*=0.004, student’s t-test). **(c)** Site modality (bimodal vs. unimodal) are not independently distributed across vegetation types (forest in green, grassland in blue, shrubland in purple, cultivated in red). Bimodal sites are more frequently associated with shrubland but less with grassland (*p*=0.019, χ^2^=9.6).

Out of the 12 factors analyzed, we selected two (one continuous and one categorical) to visualize in more detail (see Supplementary Information Fig.S3 for the rest 10 factors). Fig. 2b reveals that unimodal sites have nearly 3 times total root biomass compared to that of bimodal sites (990 *vs.* 368 g/m2). This pattern is rather unexpected yet significant despite the small sample size (*p*=0.004, student’s t-test). We also show that site vegetation cover is associated with site modality (Fig.3b; *p*=0.019, χ^2^=9.6; Methods). While the majority of bimodal sites are forested, bimodality is more frequently associated with shrublands than with grasslands.

**Figure 3.**
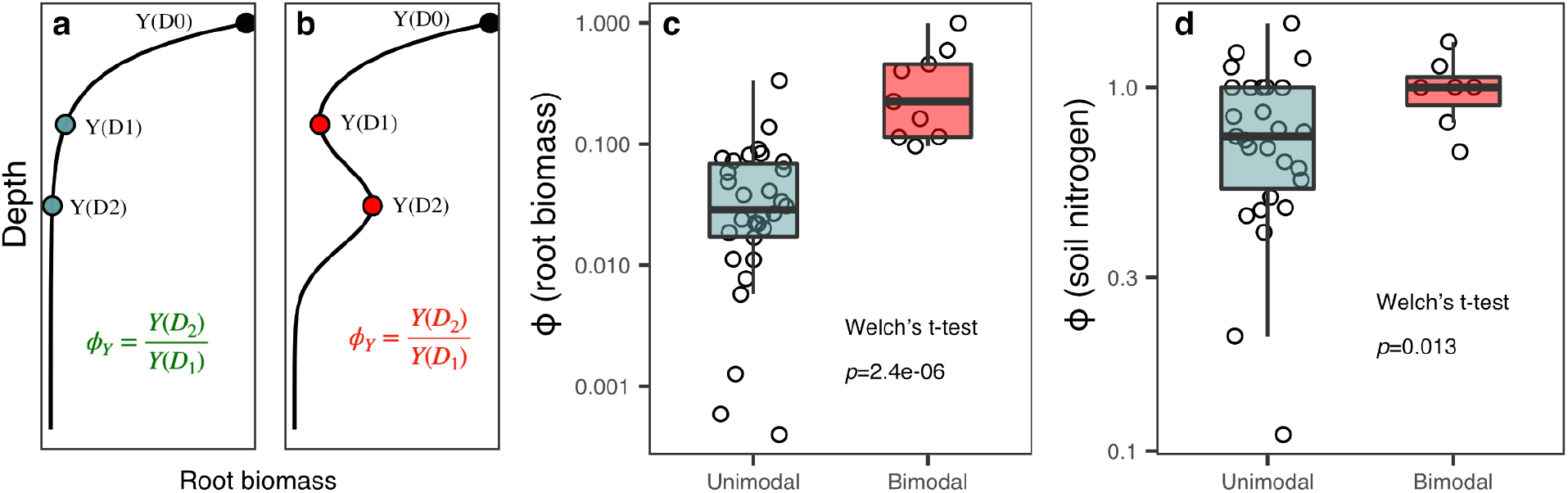
Root bimodality explained by within-site depth-level factors. (**a,b**) Illustration of a unimodal distribution (a) and a bimodal distribution (b) of property Y (root biomass in this case). Three depths derived from bimodal distribution are highlighted with filled dots: primary mode depth (D_0_), antimode depth (D_1_), and secondary mode depth (D_2_) (Methods). Using a given property value (Y) at depth D_1_ and D_2_, we can calculate ϕ as the ratio of Y(D_2_) and Y(D_1_). (**c**) Comparison of ϕ (root density) across unimodal and bimodal sites (Welch’s t-test, *p*=2.4e-06). (**d**) Comparison of ϕ (soil nitrogen concentration) across unimodal and bimodal sites (Welch’s t-test,*p*=0.013). Note that ϕ is unitless.

When we removed total root biomass from our predictive model, the resulting model has little predictive power, suggesting that no abiotic factor is a universal cause for root bimodality across all sites. This result, however, does not rule out the possibility that these factors might be important at each individual site. In fact, the finding that bimodality is strongly linked to root biomass suggests that factors that can induce low root biomass might be important for inducing root bimodality (*e.g.,* high aridity, low GPP, low soil pH), but the specific cause of low root biomass at each site can be different.

### Water- or nitrogen-driven: a depth-level analysis of bimodality

We next zoom into within-site factors along the soil depth profile (*i.e.*, depth-level) to examine whether soil nutrient concentrations and/or soil moisture may play a role in the emergence of root bimodality. To quantify the degree of bimodality in root biomass and its associated predictors, we devised a signature metric ϕ defined as the ratio of Y(D_2_) and Y(D_1_) where D_2_ is the depth of secondary root biomass mode, D_1_ the antimode (D_0_ for primary mode, see Fig.3b), and Y denotes properties such as root biomass density, soil moisture, soil nutrient content, etc. (Fig. 3a,b).

By the definition of bimodality, when we calculate ϕ for root density across 44 sites, bimodal sites have significantly higher ϕ value than unimodal sites (Fig.3c; *p*=2.4e-6, Student’s t-test). When we extend this analysis to a range of soil conditions (nutrient content, soil texture, soil moisture, etc.), our results only identify an analogous pattern in soil nitrogen concentration: bimodal sites contain significantly more nitrogen at depth D_2_ compared with the default expectation (Fig.3d).

Two alternative mechanisms might explain the presence of a nitrogen rich layer that coincides with the secondary peak of root biomass: the “nitrogen-driven hypothesis” and the “water-driven hypothesis”.

For the “nitrogen-driven hypothesis”, we speculate that the presence of nitrogen in the soil can induce fine root growth. It is not uncommon that nutrient concentrations can feature a secondary peak in deep soil due to a number of factors, such as weathering of primary rock (albeit low concentrations^42,43^), accumulation of nutrients beyond the typical root uptake zone^44^, and the presence of buried soil horizons rich in nutrients^45^. In turn, these deep nutrient anomalies can trigger the growth of fine roots which are responsible for foraging for limiting nutrients. This ability of roots responding to nutrient cues have been extensively demonstrated by physiology literature looking at root reproduction in response to localized chemical cues^46^, or in response to heterogenous nutrient patches in the soil matrix^47^.

However, the presence of deep nitrogen-rich layers could also be the result —instead of the cause— of the abundant fine roots, which might be produced to forage for other limiting resources such as water (“water-driven hypothesis”) or phosphorus. It is well known that the presence and turnover of fine roots can build up soil organic matter and consequently soil nitrogen^9,48^. It is thus possible that the nitrogen layer can be caused by the decay of water-seeking or phosphorus-seeking fine roots.

Potentially due to the limited depth-resolution of soil moisture sensors (Methods), we didn’t find definitive evidence that supports the presence of a soil moisture peak coinciding with the presence of a secondary root peak. Specifically, when we analyzed long-term dynamics of soil moisture (ϕ_soil moisture_) between unimodal vs. bimodal sites, we found no meaningful difference (Fig.S4; Methods). In evaluating whether surface drought might induce bimodality, an in-depth look at surface soil moisture (at D_0_) did not render a consistent difference between bimodal vs. unimodal sites (Fig.S5). However, in natural ecosystems, fine roots are very plastic and responsive to localized and temporally brief supply of water^49^. It is thus entirely possible that the lack of support for the “water-driven hypothesis” was simply due to low resolution (spatial and temporal) of soil moisture data.

The “nitrogen-driven hypothesis” and the “water-driven hypothesis” do not have to be mutually exclusive. After all, the process of nitrogen mineralization-microbial release of available nitrogen from soil organic matter—is often limited by water availability^50^. Moreover, it is very likely that different mechanisms (water, or different types of nutrients) might be causing bimodality at different sites. For instance, we didn’t find statistical support for phosphorus driving root bimodality, yet we cannot rule out the possibility of foraging for phosphorus being the cause of root bimodality at certain NEON sites, especially because only total rather than available phosphorus was measured at the 44 sites. Future studies with higher temporal and spatial resolution, especially controlled-experiments, might help us address these unresolved questions.

### Implications of root bimodality for subsoil nutrients and carbon

To evaluate the potential impact of root bimodality on plant resource uptake, we next analyzed the spatial distribution of soil nutrients in relation to the vertical distribution of roots. We choose to use root biomass density as a surrogate for root uptake as some biomass proliferation represents a direct plant investment for resource acquisition. In natural settings, however, the realized uptake rate often depend on nutrient availability in the soil solution^51^. But the use of root biomass density provides a good estimate of resource uptake potential, as one would expect strong correlation between root biomass and realized nutrient uptake when averaged over time and space. Alternative metrics for root uptake also include root surface area and root length (not available in our dataset), both correlate with root biomass.

When expressed relative to the abundance of fine root biomass (see Methods), relative soil nutrients appear to increase in abundance with increasing depth. For example, the relative phosphorus (P) abundance from site “YELL” (Yellowstone National Park NEON) increases with depth (Fig.4a; *r^2^* = 0.85), indicating that subsoil phosphorus is becoming relatively more abundant from the perspective of the roots.

**Figure 4.**
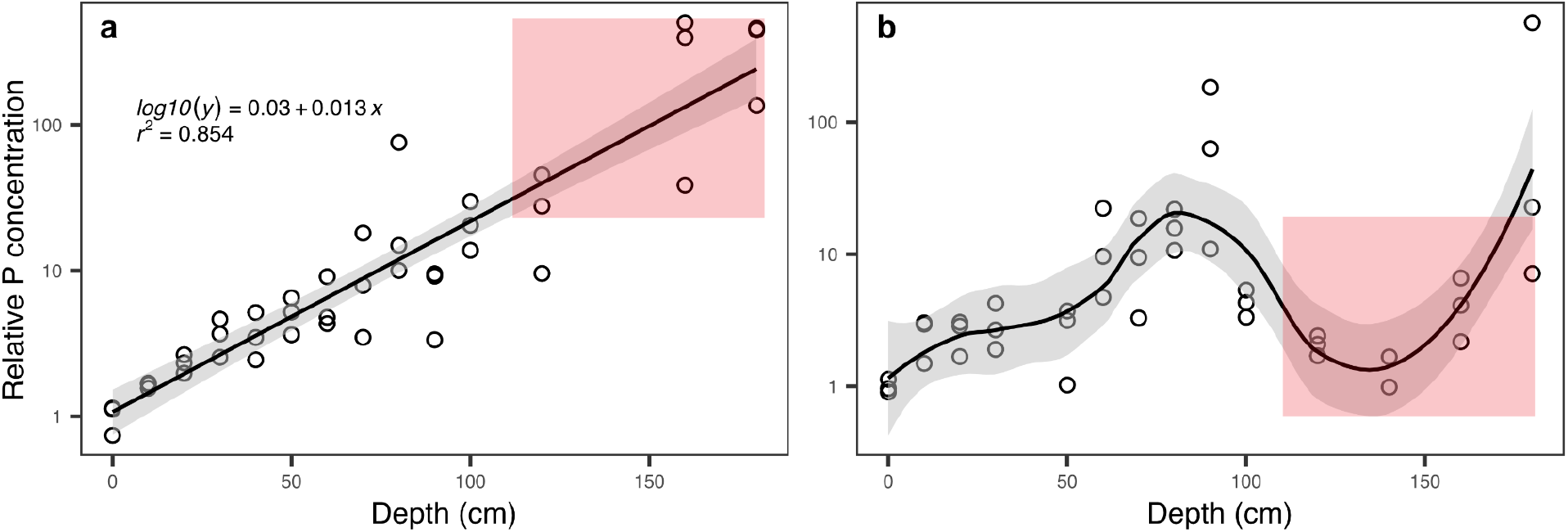
Relative nutrient concentration changes with soil depth in an unimodal (a) and a bimodal (b) site. We calculated relative nutrient concentration by dividing absolute nutrient concentration by the root abundance. For ease of comparison across sites, we rescaled relative root abundance by dividing relative abundance using topsoil value such that topsoil (Depth=0cm) relative nutrient concentration is 1(*i.e.,* dimensionless, see Methods). **a)** Using YELL as an example, both soil P content and root abundance decrease exponentially with increasing soil depth, but root abundance decays at a faster pace with depth. The resulting relative P concentration (black circle) increases exponentially with increasing soil depth (exponent = 0.013). This pattern suggests that as soil depth increases, nutrients become comparatively more abundant or underexploited in the subsoil (red area). Note that the Y axis is in logarithmic scale and the goodness of fit (*r^2^*) applies to the corresponding linear regression (log_10_(y) instead of y). **b)** Using SCBI as an example, the presence of bimodal root distribution generates a region of low relative nutrient concentration in the subsoil (red area).

We can expand the example analysis in Fig.4a to all 44 NEON sites and across four major soil nutrients: nitrogen (N), phosphorus (P), potassium (K), and calcium (Ca) (Fig.S6-S9). We can summarize the exponents across all sites for each individual nutrient, with each datapoint in Fig.S10 corresponding to the slope value of a given NEON site (for example, 0.013 in Fig.4a for P at YELL). Despite considerable variation across sites, our results clearly identify a general trend similar to what is shown in Fig.4a: *root abundance decays relatively faster than nutrients with increasing depth, leading to increasing relative nutrient abundance* (*i.e.,* positive exponent).

However, the presence of bimodal root distribution seems to disrupt the aforementioned general trend: the subsoil relative nutrient concentration in bimodal site (red area in Fig.4b, site SCBI) is dramatically lower compared to the expectation set up by the unimodal site (red area in Fig.4a, site YELL). And consequently, the goodness of regression fit is much poorer for bimodal sites across all nutrient types (Fig.S11).

Our finding that soil nutrients become relatively more abundant relative to root biomass with increasing soil depth suggests that subsoil resources might be systematically under-exploited by plant roots (Fig.4a, Fig.S10). This finding is puzzling given how frequently nitrogen^52^, phosphorus^53^, potassium^54^, and calcium^55,56^ can limit the productivity of terrestrial ecosystems, pointing to some fundamental constraints that limit plant roots to exploit subsoil. For example, many studies have argued that plants preferentially use resources from shaller depths due to: 1) the high carbon cost associated with growing deeper root systems, 2) morphologically-induced hydraulic limitations, and 3) the evaporative demand of acquiring resources from deeper depths ^44,57–61^.

However, the discovery of bimodal root distribution across a range of sites suggests otherwise (Fig.1, Fig.4b). In fact, the presence of root bimodality indicates that root depth distribution can be flexible and opportunistic: given the right condition, plants can send abundant roots to deep soil. Numerous previous studies corroborate this argument. For example, Iversen et al. (2010) reviewed the change of root distribution in response to elevated CO_2_ treatment^62^, and discovered that roots can “dig deeper” with increased carbon supply (and consequently higher nutrient demand). Dry sites or sites with high seasonality in precipitation have also shown evidence of flexible bimodal distributions wherein the depth of water uptake becomes deep in the dry season and shallow in the wet season^63–65^.

The carbon consequence of enhanced root growth in deep soil can be far-reaching^66^. On the one hand, we would expect enhanced input of new carbon due to root turnover. Deep soil carbon relies much more on root carbon input than aboveground litter input^9,11,67^. And once root carbon is deposited, the carbon decomposition rate can be 60% slower compared with the rate derived from the surface soil^11,68^, in part due to the combined effect of lower microbial community density and increased mineral protection^69–71^. Alternatively, we might also expect enhanced decomposition of old carbon induced by root production at deeper depths: physical disturbance of carbon-mineral complex^72^, impacts on the soil food web^73^, and enhanced microbial activities through priming effects^74–76^ can lead to potential destabilization of subsoil.

The implications of our finding can be profound especially against the backdrop of rapid global change. An extensive literature examines the role of increasing CO_2_ in driving an increasing degree of plant/ecosystem scale nutrient limitation^2,3,77^: plants are getting increased supply of carbon but increasingly less nutrients as more and more nutrients are locked up in plant biomass. The resulting progressive nutrient limitation has fueled the fear of a saturating land carbon sink^4^ due to the inability of the ecosystem to fix additional carbon^3,5,7^. Our analyses thus suggest plants can potentially invest an increasing amount of their photosynthates to tap into the previously under-exploited soil nutrient pool as the plant cost-benefit equation is being shifted by global warming and rising CO_2_.

### Conclusions

An improved understanding of the spatial distribution of plant roots is critical for understanding the nutritional life of land plants, and may help to predict the future trajectory of the land carbon sink. We report a surprisingly widespread occurrence of root bimodality across a broad range of ecosystems. This new understanding of the vertical distribution of plant roots could help us to better scale up point measurements to the entire soil profile, a knowledge gap especially relevant to the study of ecosystem nutrient cycles and the land carbon sink. In addition, our observation can aid in the development of the next-generation mechanistic vegetation models, which for decades have relied on relatively simplistic representation of roots (if any). But perhaps more broadly, our findings add to the growing recognition that the field of soil ecology and ecosystem ecology might have systematically overlooked dynamics and phenomena taking place in the deep soil; our results call for more research attention to this deep frontier in the face of rapid environmental change.

## Methods

### Methods

#### NEON mega-pit dataset

At each terrestrial field site, NEON collected soil at a single, temporary soil pit (*i.e.*, the megapit). The pit was selected to be near the instrumented NEON tower and soil sensors. In each pit, three soil profiles were sampled and described up to 2m deep (see SI Fig.1 for exact profile depth across sites). Depth-specific chemical and physical properties of soil samples and root samples from these megapit profiles were analyzed in collaboration with the US Department of Agriculture (USDA), Natural Resources Conservation Service Kellogg Soil Survey Laboratory(NRCS-KSSL). We accessed via the neonUtilities R package^78^ the following NEON data products: Root biomass and chemistry, Megapit (DP1.10066.001)^79^ and Soil physical and chemical properties, Megapit (DP1.00096.001)^80^. All NEON datasets used in our analyses are summarized in supplementary information Table.S2.

#### Soil moisture sensor data

We downloaded soil moisture time series from all sites with data ranging between 2018-Sep and 2021-Aug (3 year span). Soil sensor measures soil water content/ion content across a 10cm vertical measurement zone (5cm above and below). Sensor depth represents the center of the measurement zone, at 6cm, 16cm, 26cm, 56cm, 86cm, 116cm, 146cm, 176cm (exact depth varies across sites, exact depths can be found in our data deposit). To ensure data coverage, multiple soil moisture sensors were installed at each depth across different profiles within each site. Two time resolutions were provided from the soil moisture data product: 1-min resolution vs. 30-min. Given plant rooting systems are unlikely to be sensitive to 1min-level soil moisture variation, we used soil moisture sensor data that was averaged at 30mins (DP1.00094.001)^81^.

#### Determining site modality

We first selected live fine roots from all profiles across all sites (3 profiles per site) and merged profile level data into a site level mean root density at each depth. We then translated biomass data (double precision) into frequency of counts (integer) in order to process the modality of biomass depth distribution. Using R package “multimode”, we can derive the number, magnitude, and location of mode/modes in a distribution^82,83^. Given the root sampling depth is not continuous according to NEON sampling procedure, we used a bandwidth of 15cm for kernel density estimation such that the algorithm would not pick up each sampling depth as a distinct mode. We also defined a threshold for classifying bimodal distribution: the secondary peak has to be equal or larger to 10% of the magnitude of the primary peak. Increasing this cutoff value will select for sites with more prominent secondary peaks but leaves few sites qualifying. In theory, our approach allows for detection of more than 2 modes, but in practice, a maximum of 2 modes was detected (especially after applying the threshold for mode magnitude). For bimodal distribution, depth locations of each mode and antimode are recorded as (D_0_, D_1_, D_2_) for primary mode, antimode, secondary mode respectively.

#### Calculating ϕ

We devised a profile-level metric ϕ that describes the ratio of property Y(D_2_) and Y(D_1_) where D_2_ identify the depth of secondary root biomass mode, D_1_ the antimode (D_0_ for primary mode), and Y denote properties such as root biomass density, soil nutrient content, soil moisture, etc (Fig.3AB). For unimodal sites (n=35) where antimode depth D_1_ and secondary model depth D_2_ is not available, we prescribe D_1_=70cm, D2=100cm that is consistent with the mean value from across all bimodal sites (n=9).

For root biomass and soil nutrient content that is only sampled once, we derive a single ϕ value per site. For soil moisture data however, we derive a time series of ϕ for soil moisture at each profile within each site. The use of time series (instead one time point) would allow us to examine the entire dynamics of soil moisture instead of just the mean or median, as the root status we observe (at a single time point) might be a response to a past short-period of soil moisture stress or incentive. For each time series, we visualize the full frequency distribution of soil moisture and subsequently calculate a list of distribution metrics: mean, median, max, min, standard deviation, coefficient of variance, 75th quantile, 25th quantile, interquartile range.

The depth resolution of our soil moisture data resolution (roughly, 6cm, 16cm, 26cm, 56cm, 86cm, 116cm, 146cm, 176cm) might not be ideal to distinguish D_1_ and D_2_ for certain sites. To compensate for this low depth resolution of soil moisture, we additionally examined the soil moisture dynamics of the surface sensor (6cm) to evaluate whether the presence of bimodality is a result of surface drought.

#### Relative nutrient concentration

To calculate relative nutrient concentration in Fig.4, we first transformed the original nutrient concentration (N, P, K, or Ca) by dividing them using the root biomass density (mg/cm3) across all profiles of all 44 sites. For ease of cross-site comparison, we then rescaled the resulting relative nutrient concentration using relative nutrient concentration at surface depth. The rescaled relative nutrient concentration is thus dimensionless and the surface soil has an average value 1.

#### Statistics

We developed a classification model based random forest algorithm (Fig.2a; R package randomForest). Given the unbalanced nature of our dataset (9 bimodal vs. 35 unimodal sites), we need to make sure that we predict the minority class (bimodal sites) equally well as the majority class (unimodal sites). The default behavior of random forest algorithms will prioritize the prediction accuracy of the majority class (thus maximizing overall accuracy). As a result, we modified the default voting rule such that the algorithm weighs more on the prediction accuracy of the minority class. We next searched the best performing model (highest bimodality prediction accuracy) by finding the best parameter combination across a grid of conditions (num.trees = 500, mtry = 6, min.node.size = 3,sample.fraction = 0.3, seed = 123). The best performing model after removing profile root biomass has the following parameter combination: num.trees = 500, mtry = 6, min.node.size = 3,sample.fraction = 0.3, seed = 123. The feature importance score shown in Fig.2a is based on mean decrease in Gini coefficient, a measure of out-of-bag cross validated predictions. We sequentially dropped factors with the lowest Gini coefficient (such as temperature range, dry season precipitation, precipitation seasonality, etc.), when trimming the model would enhance the overall performance of the model. We used t-test to examine the feature differences between bimodal and unimodal sites (Fig.2b, Fig.S3). We first visualize the distribution of each feature splitted by modality and log_10_ transforms the data to conform with normality. Then we used the F test to test equal variance between the unimodal and bimodal group to determine the use of Student’s (equal variance) or Welch’s t-test (unequal variance). We used the χ^2^ test of independence to test the association between site modality and site vegetation cover (Fig.2c). Due to the small sample size (N=44), we used the Pearson’s Chi-squared test with simulated p-value (R base function *chisq. test,* simulate.p.value = true, 2000 replicates). We used linear regression to fit the relationship between rescaled relative nutrient abundance and depth (Fig.4a,b). Rescaled relative nutrient abundance is log_10_ transformed in all regression analyses, consistent with our visualization. All statistical analyses were performed using the R platform (R version 4.0.5).

## Code availability

R scripts are deposited together with associated data on figshare (see data availability) and available from the corresponding author upon request.

## Data availability

All data will be deposited at figshare (DOI: 10.6084/m9.figshare.19525303).

## Acknowledgments

This work was supported by the Santa Fe Institute Omidyar Fellowship to M.L. We thank resident faculty members and postdocs at the Santa Fe Institute and members of the Jackson lab for helpful comments on the manuscript. The National Ecological Observatory Network (NEON) is a program sponsored by the National Science Foundation and operated under cooperative agreement by Battelle. This material is based in part upon work supported by the National Science Foundation through the NEON Program.

## Author contributions

M.L. and S.W. conceived and designed research, M.L. performed analyses with input from S.W., R.B.J., A.M, and S.J.T., M.L. wrote the paper and all authors contributed significantly to writing.

## Additional Information

Correspondence and requests for materials should be addressed to M.L.

## Supplementary Information Table of Content

**Supplementary Information Table S1**| NEON site description.

**Supplementary Information Table S2**| NEON data products used in the analyses.

**Supplementary Information Figure S1**| Maximum profile depth across all 44 sites.

**Supplementary Information Figure S2**| Root biomass depth distribution across all 44 NEON sites.

**Supplementary Information Figure S3**| Feature comparison between bimodal and unimodal sites.

**Supplementary Information Figure S4**| Frequency distribution of ϕ_soil moisture_ across all NEON sites.

**Supplementary Information Figure S5**| Frequency distribution of surface soil moisture.

**Supplementary Information Figure S6**| Depth distribution of rescaled relative nitrogen abundance (N) across 44 NEON sites.

**Supplementary Information Figure S7**| Depth distribution of rescaled relative nitrogen abundance (P) across 44 NEON sites.

**Supplementary Information Figure S8**| Depth distribution of rescaled relative nitrogen abundance (K) across 44 NEON sites.

**Supplementary Information Figure S9**| Depth distribution of rescaled relative nitrogen abundance (Ca) across 44 NEON sites.

**Supplementary Information Figure S10**| Frequency distribution of exponents from Fig.S6-9.

**Supplementary Information Figure S11**| Summary of goodness of fit for linear regression in Fig.S6-9.

**Table S1.** NEON site description.

| Site | Lat | Long | Elev | MaxDepth | RootBio | Size | Domain | MDepth |
| --- | --- | --- | --- | --- | --- | --- | --- | --- |
| BARR | 71.28 | -156.65 | 7 | 90 | 5823.52 | <=2mm | D18 | NA |
| BONA | 65.15 | -147.50 | 186 | 168 | 800.68 | <=2mm | D19 | NA |
| DEJU | 63.88 | -145.75 | 519 | 90 | 2271.41 | <=4mm | D19 | NA |
| HEAL | 63.88 | -149.22 | 670 | 85 | 5409.78 | <=2mm | D19 | NA |
| DCFS | 47.16 | -99.11 | 571 | 140 | 650.91 | <=2mm | D09 | NA |
| WOOD | 47.13 | -99.24 | 588 | 150 | 736.35 | <=2mm | D09 | NA |
| NOGP | 46.77 | -100.92 | 588 | 200 | 1543.99 | <=2mm | D09 | NA |
| UNDE | 46.14 | -89.32 | 533 | 200 | 295.91 | <=2mm | D05 | 88.62 |
| WREF | 45.82 | -121.96 | 383 | 200 | 4502.03 | <=2mm | D16 | NA |
| ABBY | 45.76 | -122.33 | 367.6 | 200 | 2452.61 | <=4mm | D16 | NA |
| STEI | 45.51 | -89.59 | 484 | 200 | 746.02 | <=4mm | D05 | NA |
| TREE | 45.49 | -89.59 | 481 | 200 | 3529.87 | <=4mm | D05 | NA |
| YELL | 44.96 | -110.54 | 2132 | 200 | 1058.99 | <=2mm | D12 | NA |
| BART | 44.07 | -71.29 | 284 | 160 | 1249.64 | <=2mm | D01 | NA |
| HARV | 42.54 | -72.18 | 373 | 130 | 1630.89 | <=2mm | D01 | NA |
| CPER | 40.81 | -104.74 | 1548 | 160 | 605.59 | <=2mm | D10 | NA |
| STER | 40.46 | -103.03 | 1369 | 160 | 42.52 | <=4mm | D10 | 62.49 |
| RMNP | 40.28 | -105.55 | 2758 | 120 | 402.51 | <=2mm | D10 | NA |
| ONAQ | 40.18 | -112.45 | 1662 | 200 | 853.04 | <=4mm | D15 | 64.29 |
| NIWO | 40.05 | -105.58 | 3474 | 140 | 2052.66 | <=4mm | D13 | NA |
| KONZ | 39.10 | -96.56 | 414 | 200 | 1416.41 | <=4mm | D06 | NA |
| BLAN | 39.06 | -78.07 | 177 | 180 | 446.99 | <=2mm | D02 | NA |
| UKFS | 39.04 | -95.20 | 310 | 200 | 994.06 | <=4mm | D06 | NA |
| SCBI | 38.89 | -78.14 | 410 | 200 | 435.82 | <=2mm | D02 | 138.82 |
| SERC | 38.89 | -76.56 | 27.4 | 180 | 525.63 | <=4mm | D02 | NA |
| MOAB | 38.25 | -109.39 | 1805 | 180 | 521.44 | <=2mm | D13 | NA |
| MLBS | 37.38 | -80.52 | 1177 | 120 | 556.08 | <=2mm | D07 | NA |
| SJER | 37.11 | -119.73 | 374 | 180 | 338.97 | <=2mm | D17 | NA |
| SOAP | 37.03 | -119.26 | 1204 | 200 | 1799.7 | <=2mm | D17 | NA |
| TEAK | 37.01 | -119.01 | 2145 | 200 | 1228.37 | <=2mm | D17 | 91.06 |
| GRSM | 35.69 | -83.50 | 599 | 200 | 1480.22 | <=4mm | D07 | NA |
| ORNL | 35.58 | -84.17 | 366 | 180 | 912.32 | <=2mm | D07 | NA |
| OAES | 35.41 | -99.06 | 515 | 40 | 115.58 | <=2mm | D11 | NA |
| CLBJ | 33.40 | -97.57 | 288 | 160 | 999.73 | <=2mm | D11 | NA |
| TALL | 32.95 | -87.39 | 154 | 200 | 535.63 | <=2mm | D08 | NA |
| JORN | 32.59 | -106.84 | 1327 | 180 | 262.95 | <=2mm | D14 | NA |
| DELA | 32.54 | -87.80 | 18 | 160 | 594.11 | <=2mm | D08 | 89.16 |
| SRER | 31.91 | -110.84 | 996 | 180 | 198.22 | <=2mm | D14 | 88.83 |
| LENO | 31.85 | -88.16 | 8 | 200 | 835.1 | <=4mm | D08 | NA |
| JERC | 31.20 | -84.47 | 49 | 180 | 431.41 | <=2mm | D03 | 139.65 |
| OSBS | 29.69 | -81.99 | 55 | 170 | 620.73 | <=2mm | D03 | NA |
| DSNY | 28.13 | -81.43 | 25 | 120 | 559.64 | <=2mm | D03 | NA |
| LAJA | 18.02 | -67.08 | 40 | 180 | 422.08 | <=4mm | D04 | 139.23 |
| GUAN | 17.97 | -66.87 | 133 | 200 | 1408.03 | <=4mm | D04 | NA |

“Elev”, elevation in meter; “MaxDepth”, maximum profile depth in cm; “RootBio”, root biomass in g/m<sup>2</sup>; “Size”, size cutoff for fine roots; “Domain”, NEON assigned ecological domain; “MDepth” stands for secondary mode depth in cm. A full-size site description is included in the figshare deposit and available through NEON (https://www.neonscience.org/field-sites/explore-field-sites)

**Table S2.** NEON data products used in the analyses.

| **Data product** | **Data product name** | **DOI** | **Site ID** | **Date range** |
| --- | --- | --- | --- | --- |
| DP1.10066.001 | Root biomass and chemistry (Megapit) | https://doi.org/10.48443/xmbe-7b55 | Table S1 | 2012-2018 |
| DP1.00096.001 | Soil physical and chemical properties (Megapit) | https://doi.org/10.48443/10dn-8031 | Table S1 | 2012-2018 |
| DP1.00094.001 | Soil water content and water salinity | https://doi.org/10.48443/ghry-qw46 | Table S1 | 2018-2021 |

**Supplementary Information Figure S1.**
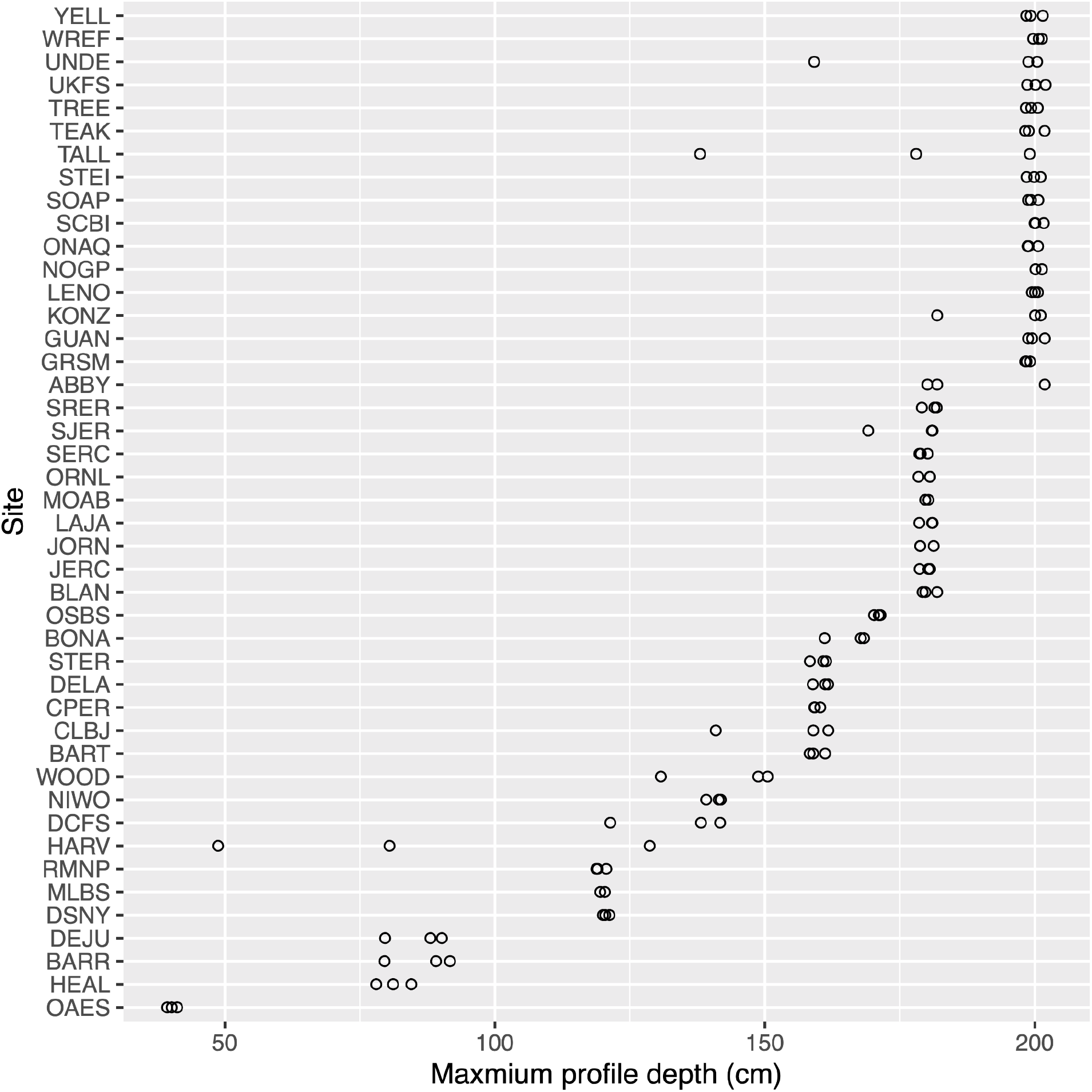
Maximum profile depth across all 44 sites. A horizontal jitter of 2 cm was applied to visualize overlapping data points. Corresponding site description for each site ID can be found on the official NEON website (https://www.neonscience.org/field-sites/explore-field-sites).

**Supplementary Information Figure S2.**
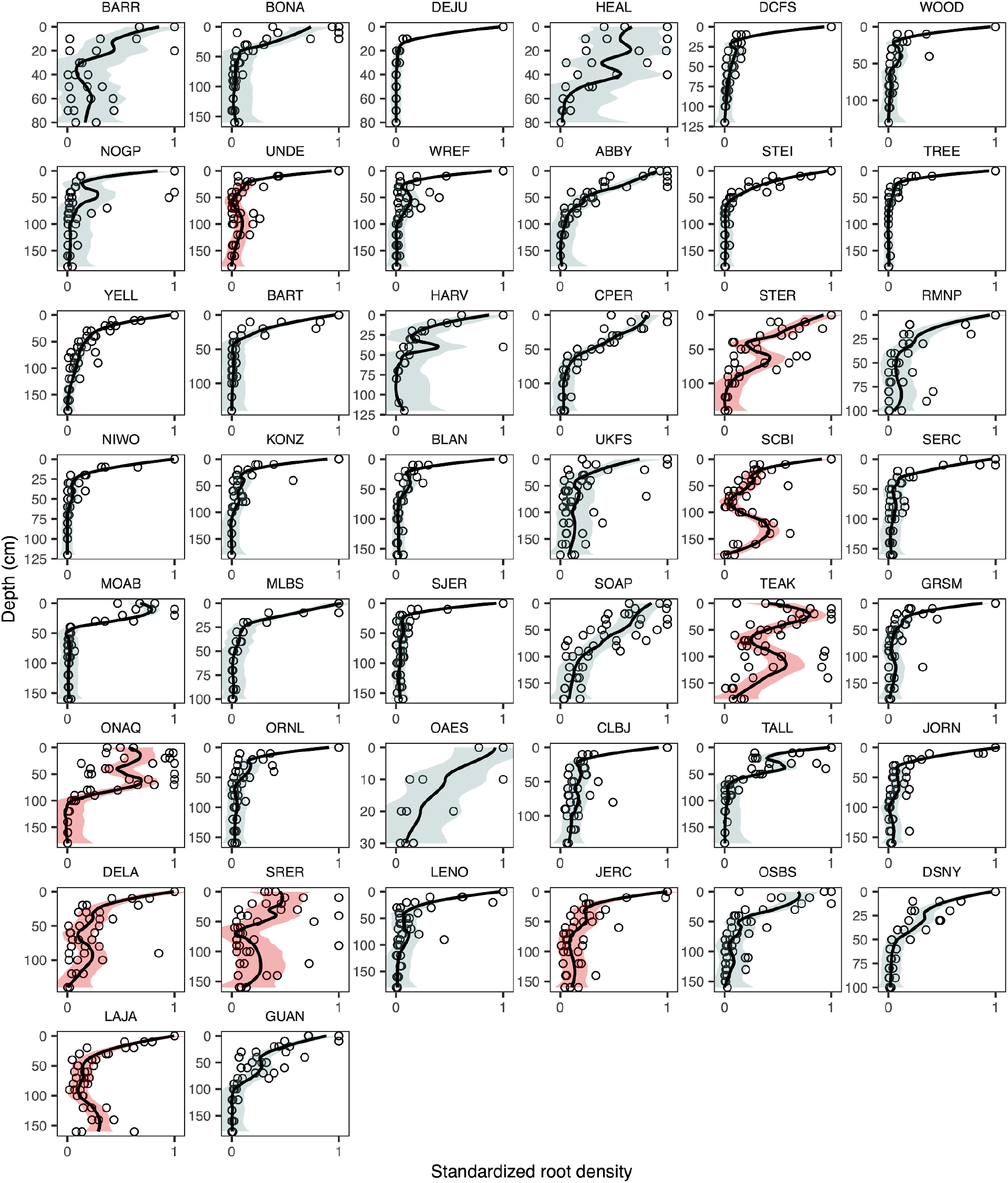
Root biomass depth distribution across all 44 NEON sites. For clearer visual representation, we standardized root biomass density by dividing all values using the maximum such that standardized root density is bounded within [0,1].

**Supplementary Information Figure S3.**
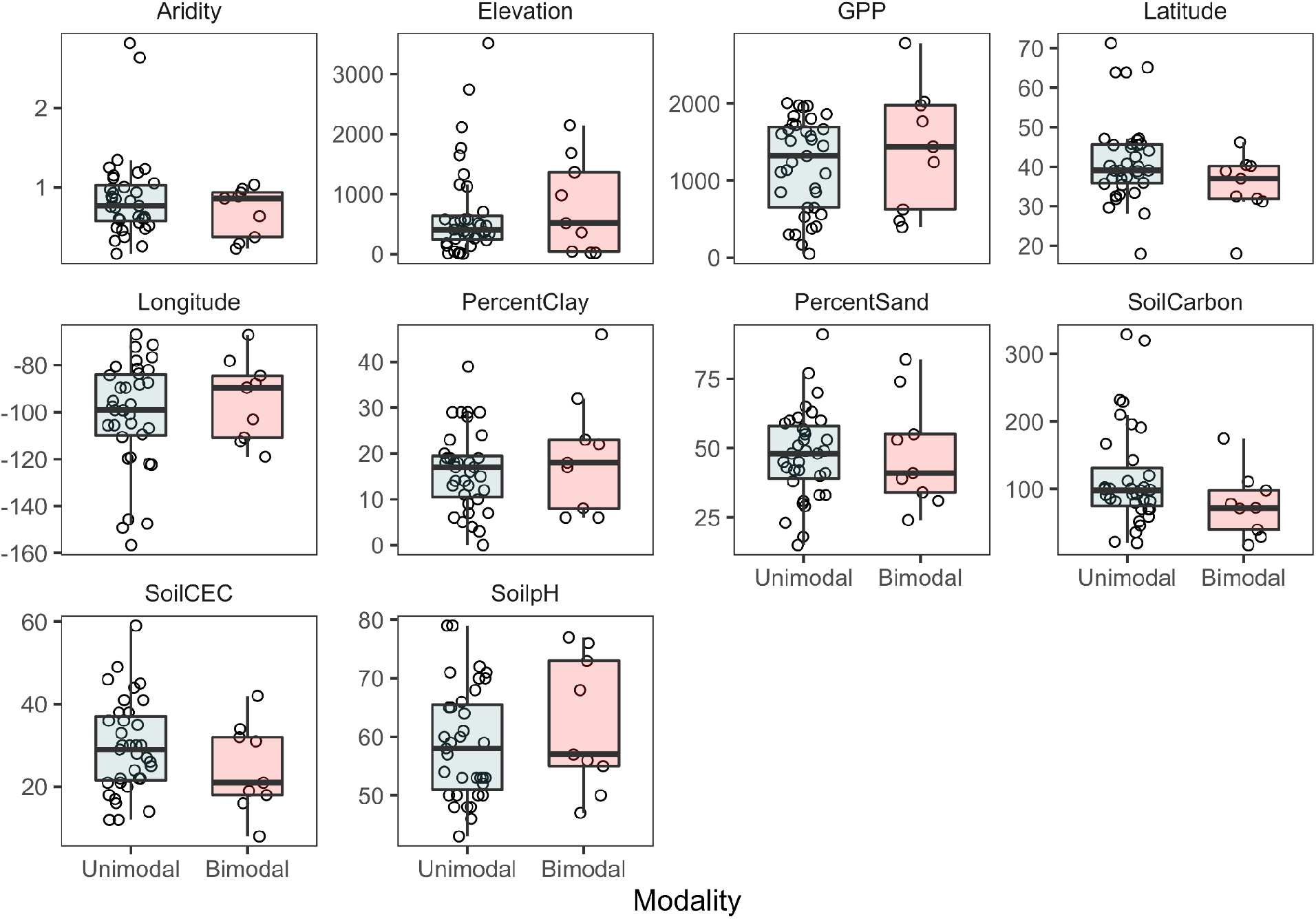
Feature comparison between bimodal and unimodal sites. We didn’t observe significant differences between unimodal and bimodal sites for these features.

**Supplementary Information Figure S4.**
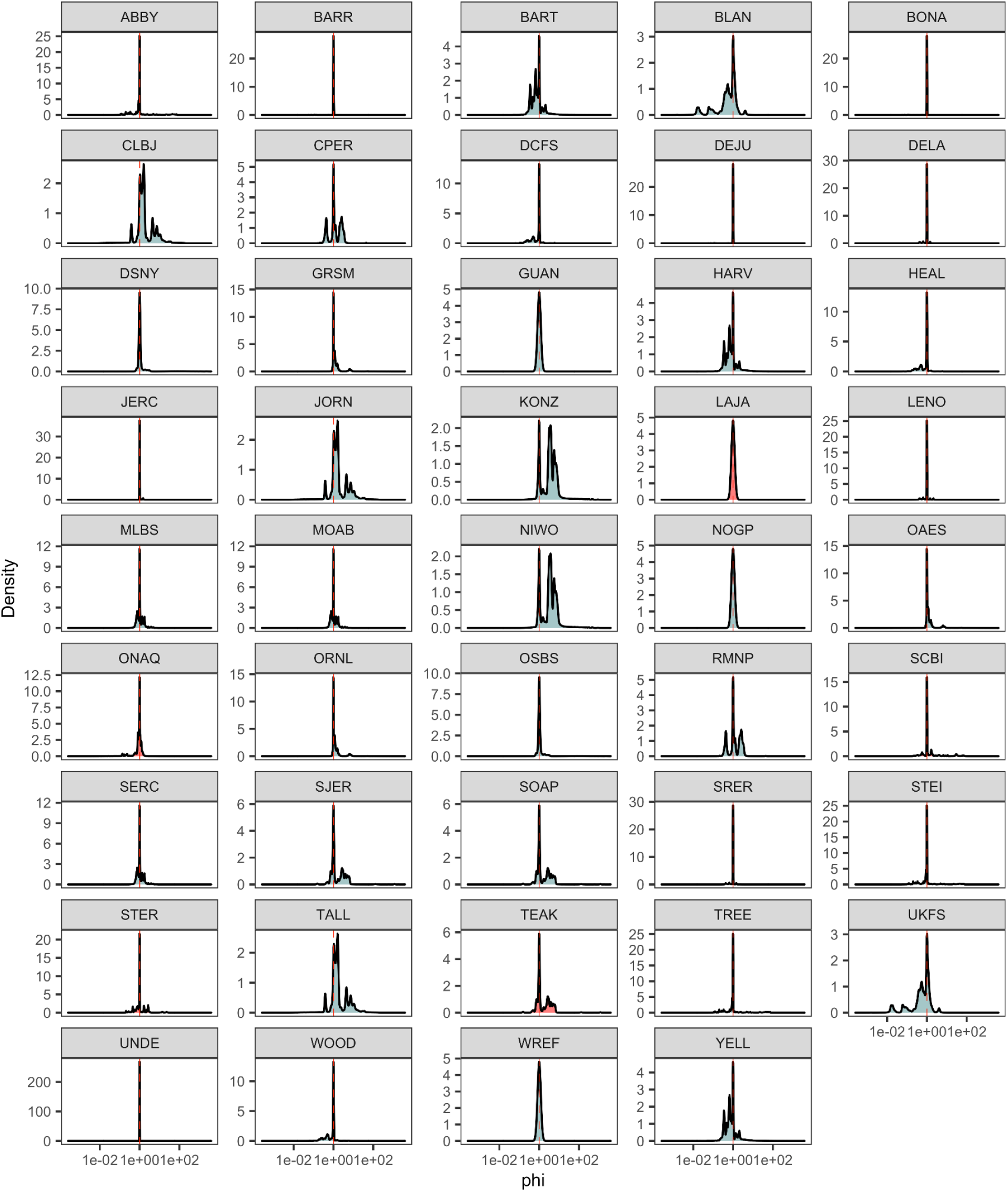
Frequency distribution of ϕ_soil moisture_ across all NEON sites. ϕ_soil moisture_ is derived by dividing soil moisture value at D_2_ (secondary mode depth, Fig.3b; Methods) using soil moisture at D_1_ (antimode depth, Fig.3b; Methods). Potentially due to the low depth resolution of soil moisture data, the most frequent ϕ_soil moisture_ is 1, indicating soil moisture at D_1_ and D_2_ is the same for most of the time.

**Supplementary Information Figure S5.**
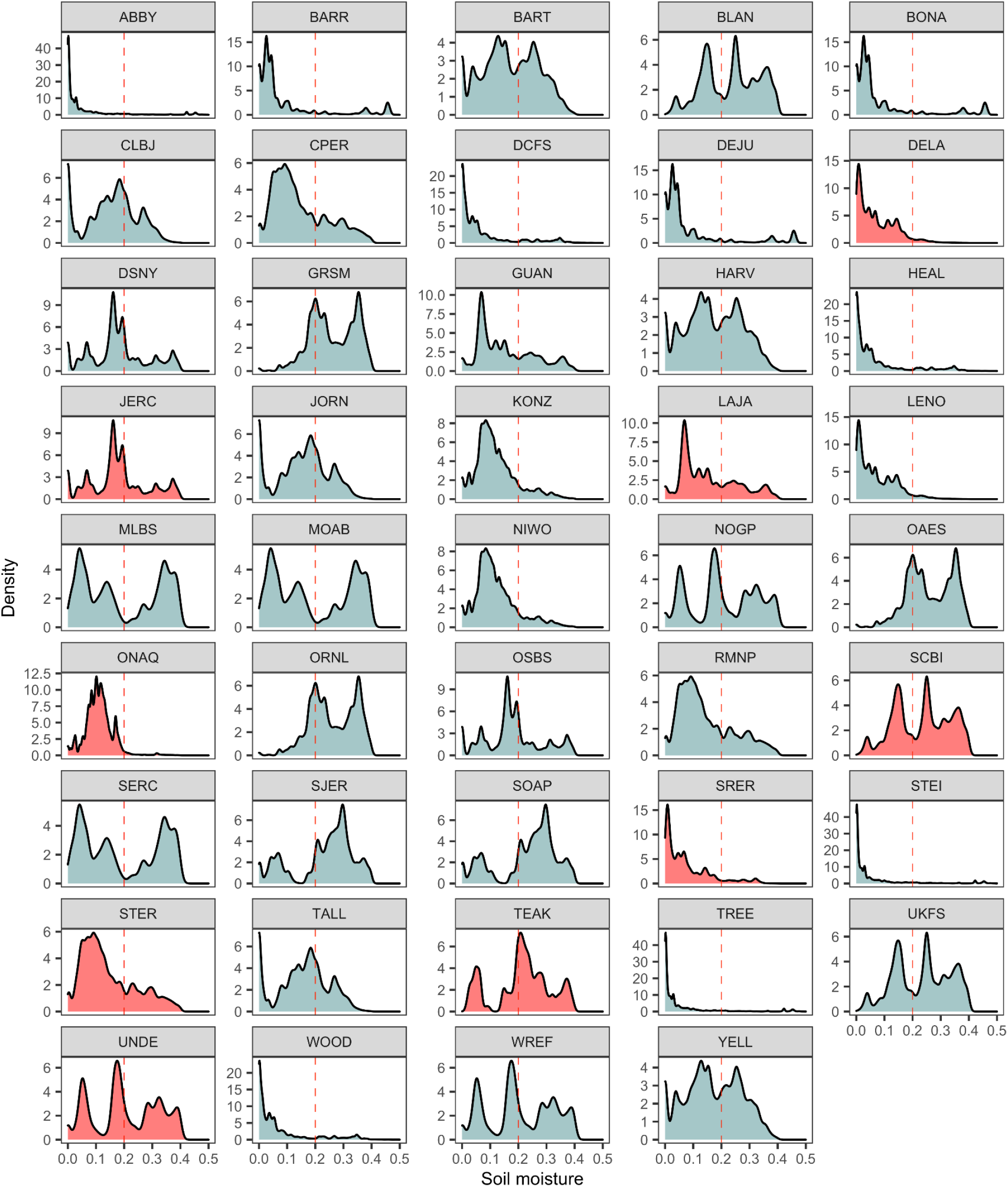
Frequency distribution of surface soil moisture. We used soil moisture at depth D_0_ (primary mode depth, Fig.3b; Methods). Unimodal sites in blue and bimodal sites in red.

**Supplementary Information Figure S6.**
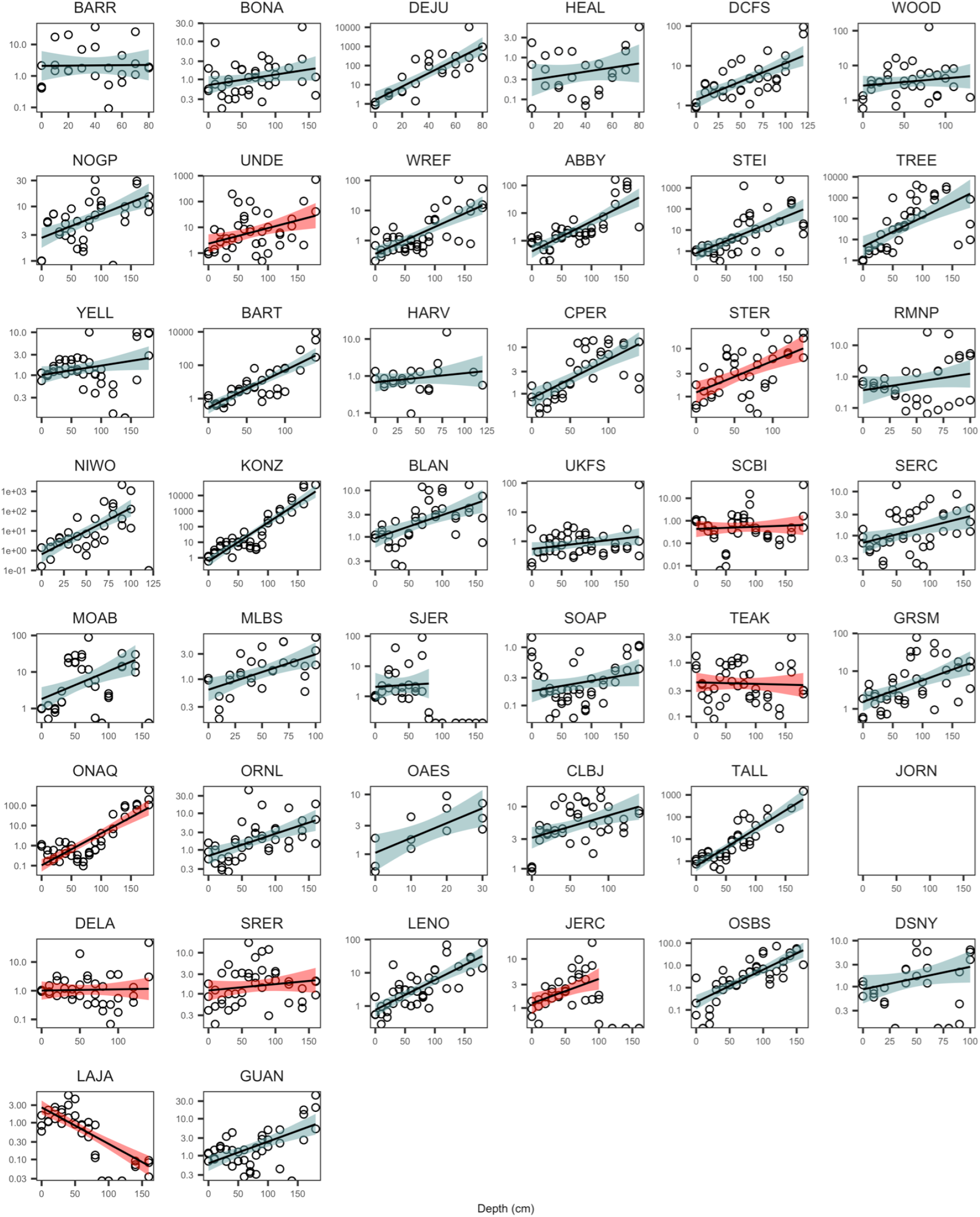
Depth distribution of rescaled relative nitrogen abundance (N) across 44 NEON sites. We calculated relative nutrient concentration by dividing absolute nutrient concentration with root abundance. For ease of cross-site comparison, we then rescaled the resulting relative nutrient concentration using relative nutrient concentration at surface depth. The rescaled relative nutrient concentration is thus dimensionless and the surface soil has an average relative concentration of value 1. Unimodal sites in blue and bimodal sites in red.

**Supplementary Information Figure S7.**
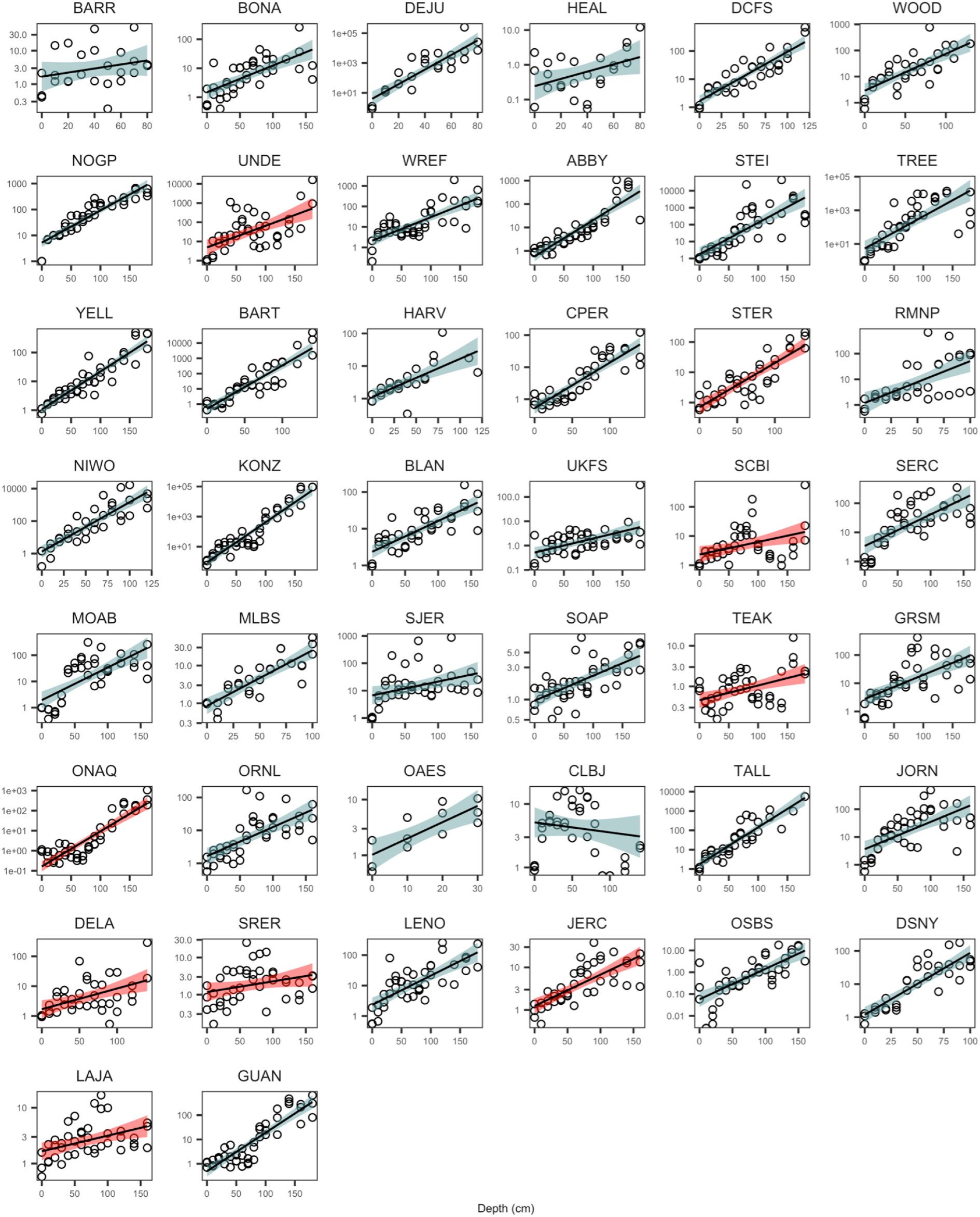
Depth distribution of relative phosphorus abundance (P) across 44 NEON sites. We calculated relative nutrient concentration by dividing absolute nutrient concentration with root abundance. For ease of cross-site comparison, we then rescaled the resulting relative nutrient concentration using relative nutrient concentration at surface depth. The rescaled relative nutrient concentration is thus dimensionless and the surface soil has an average relative concentration of value 1. Unimodal sites in blue and bimodal sites in red.

**Supplementary Information Figure S8.**
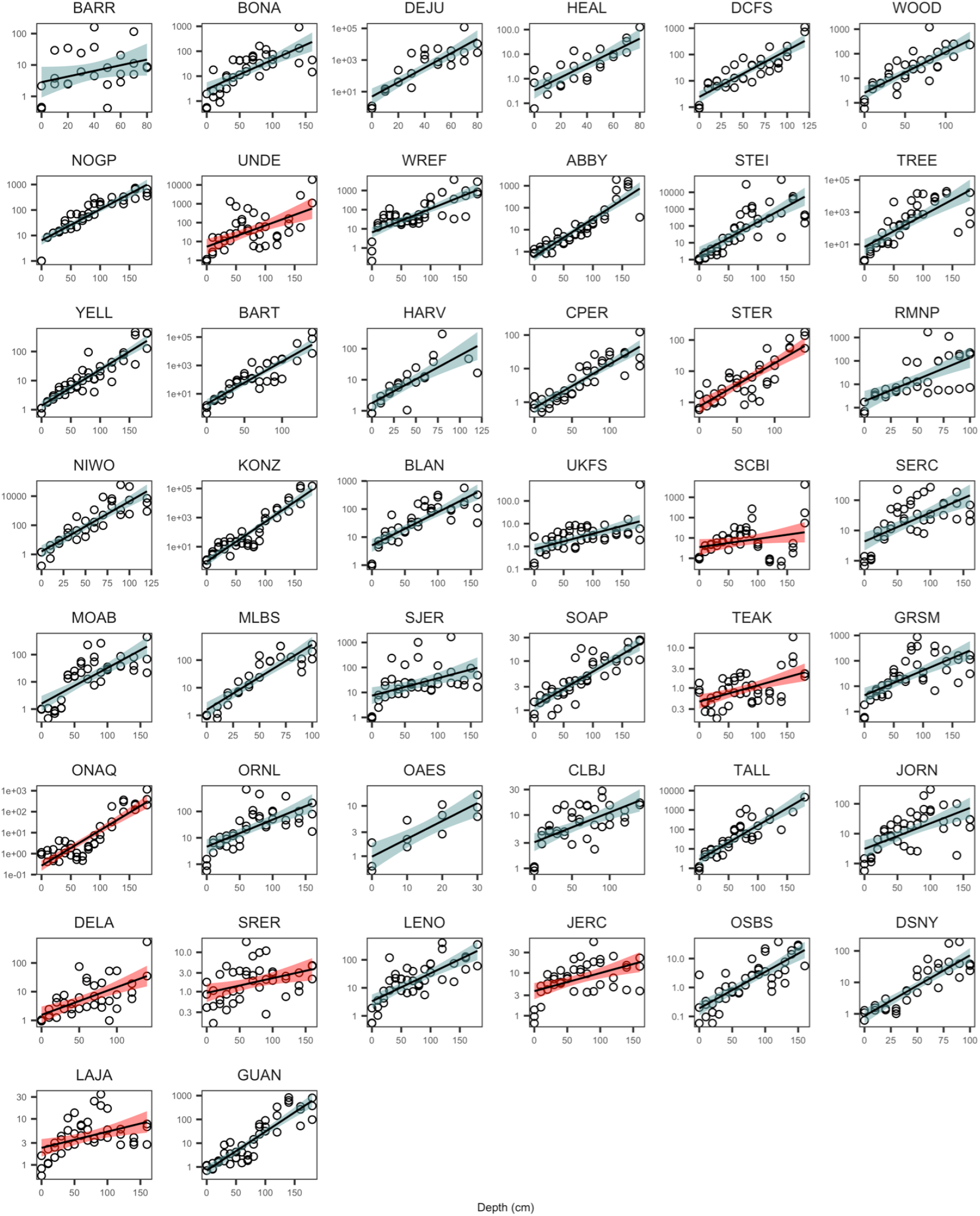
Depth distribution of relative potassium abundance (K) across 44 NEON sites. We calculated relative nutrient concentration by dividing absolute nutrient concentration with root abundance. For ease of cross-site comparison, we then rescaled the resulting relative nutrient concentration using relative nutrient concentration at surface depth. The rescaled relative nutrient concentration is thus dimensionless and the surface soil has an average relative concentration of value 1. Unimodal sites in blue and bimodal sites in red.

**Supplementary Information Figure S9.**
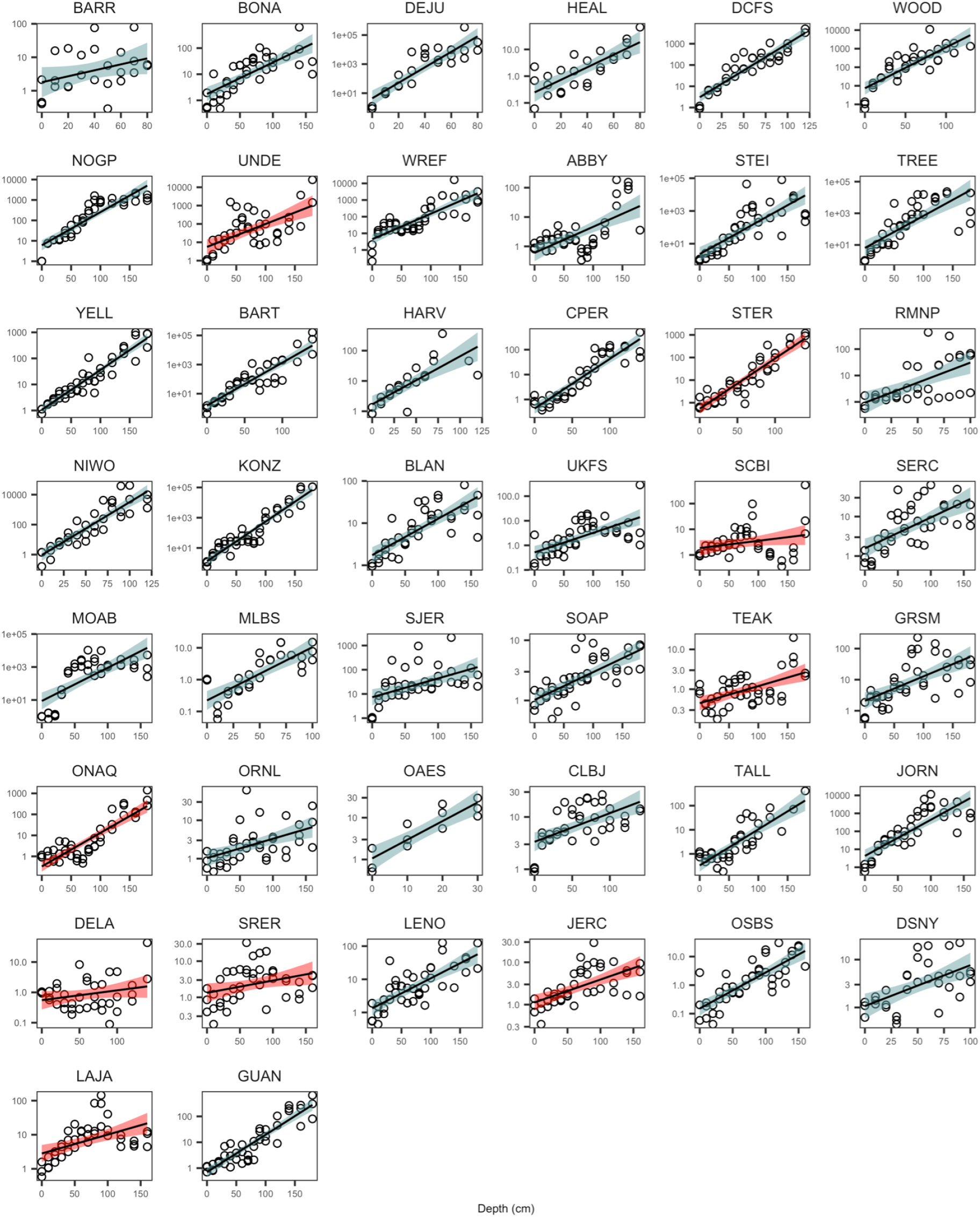
Depth distribution of relative calcium abundance (Ca) across 44 NEON sites. We calculated relative nutrient concentration by dividing absolute nutrient concentration with root abundance. For ease of cross-site comparison, we then rescaled the resulting relative nutrient concentration using relative nutrient concentration at surface depth. The rescaled relative nutrient concentration is thus dimensionless and the surface soil has an average relative concentration of value 1. Unimodal sites in blue and bimodal sites in red.

**Supplementary Information Figure S10.**
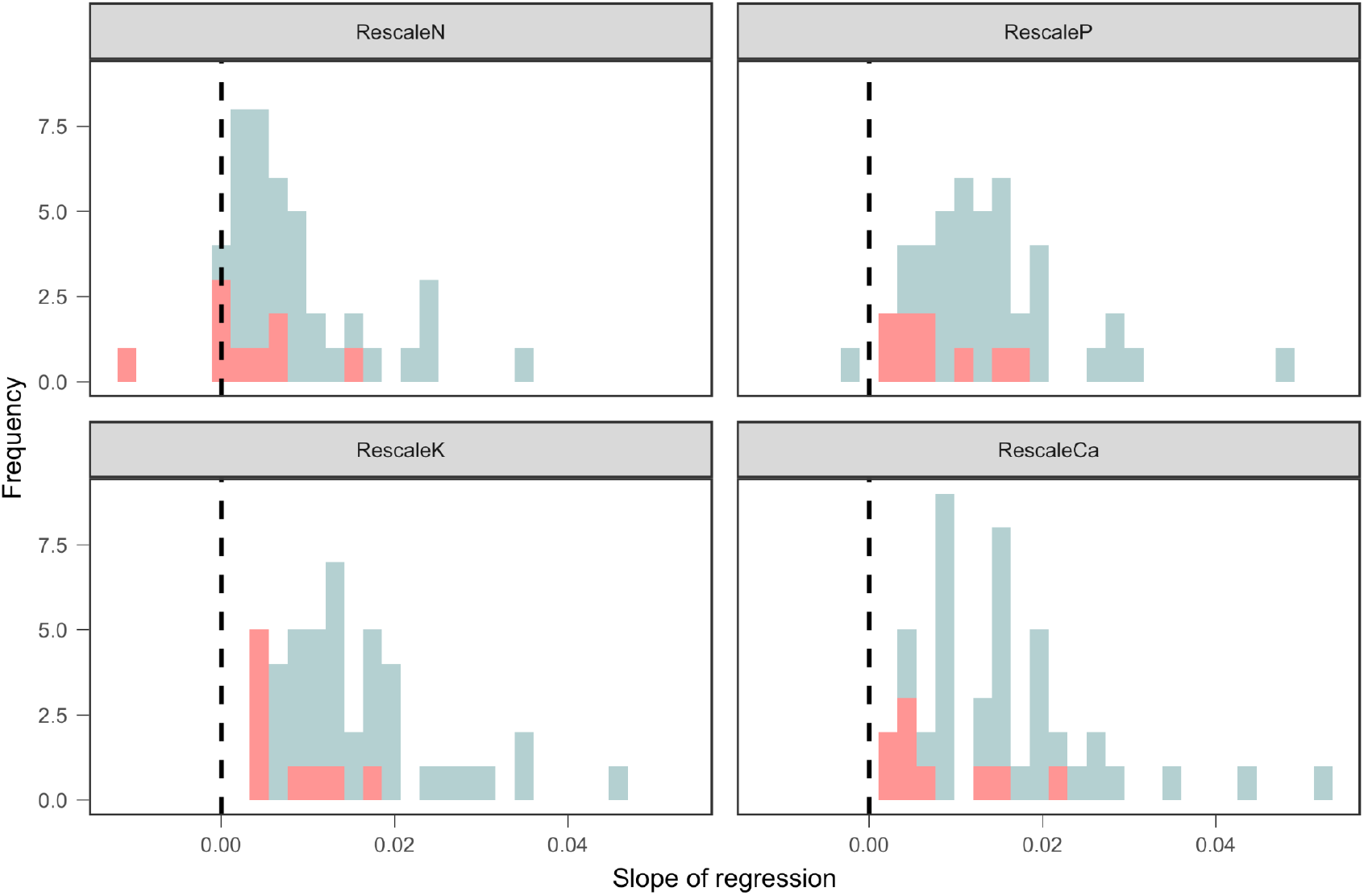
Frequency distribution of exponents from Fig.S6-9. A positive exponent (i.e., a positive slope on a logarithmic scale) suggests that nutrients concentration increases with depth relative to the abundance of roots.

**Supplementary Figure S11.**
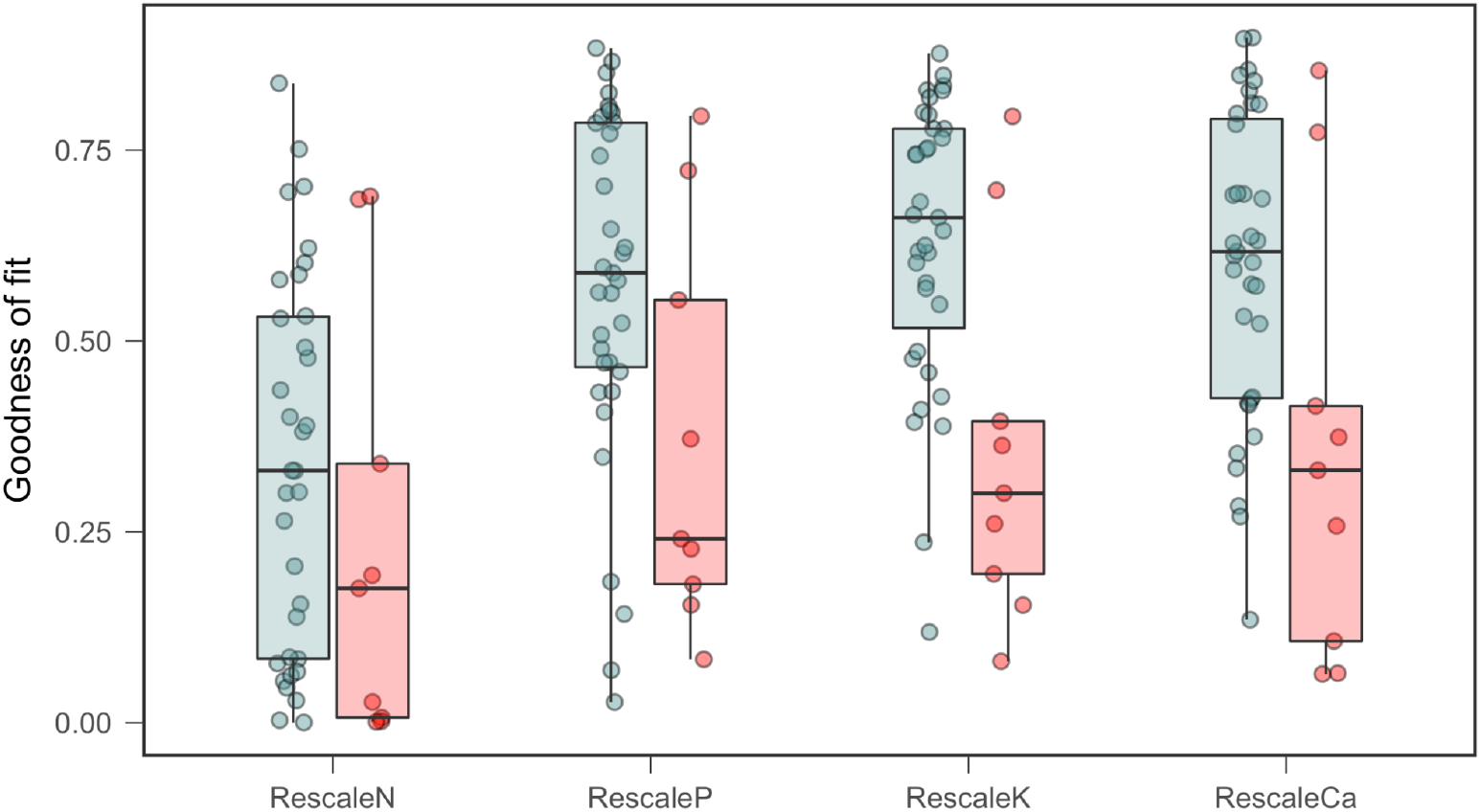
Summary of goodness of fit for linear regression in Fig.S6-9. Bimodal sites tend to have worse fit compared to unimodal sites.

